# Electrospray Surface Charge Describes Protein Molecular Motion

**DOI:** 10.1101/571091

**Authors:** Rod Chalk, Oktawia Borkowska, Kamal Abdul Azeez, Stephanie Oerum, Petra Born, Opher Gileadi, Nicola Burgess-Brown

**Affiliations:** Structural Genomics Consortium, Oxford University, UK; Max Planck Institute, Dresden, Germany

**Keywords:** Electrospray ionisation, molecular motion, mass/charge, native mass spectrometry, protein structure

## Abstract

The electrospray mass/charge distribution for a protein is an instantaneous measurement of surface areas during the transition from liquid to gas phase. Protonation is dependent upon surface area and surface area is related to protein folding under all conditions. M/z distributions for proteins and protein analogues all with different degrees of entropy were compared. Rigid dipeptide self-assemblies of any size possess a single m/z, whereas protein always displays multiple m/z distributions. Native proteins have narrow, defined m/z distributions, while denatured proteins and synthetic homopolypetides possess the widest possible m/z distributions. These observations are consistent with dynamic changes in surface area resulting from molecular motion.

## Introduction

Protein structure is not fixed, but continually changing as bonds rotate and stretch within defined stability constraints due to thermal motion. This motion is functionally essential in enabling enzyme catalysis [1] or other molecular interactions. Optimal drug design requires an understanding of transitional conformational states. Unlike NMR, X-ray crystallography and electron microscopy only provide a time-averaged structure, hence molecular dynamic simulation is generally used to explore protein molecular motion[2].

Electrospray ionisation (ESI) is the favoured soft ionisation mechanism for protein analysis by mass spectrometry [3] and analysis of proteins under native conditions has been established for over two decades eg. [4, 5]. Despite its importance, ESI is poorly understood. It is generally assumed that charge state in solution will be a guide to charge state in the gas phase, but there is no evidence to support this. Carbeck *et al.* [6] found no correlation and suggested that whereas ionisation in solution depended upon the chemical properties of a protein, electrospray ionisation depended on its physical properties, namely surface area. Neutral species with no ionisable groups readily form electrospray ions in both positive and negative ion modes, as do highly negatively charged species such as nucleic acids and glycans. All proteins generate multiple m/z values in electrospray ionisation under all conditions. Generalizations of this kind are seldom possible in biology, but we have analysed over 10,000 proteins in our laboratory and have always observed this [7]. Indeed, the process of deconvolution which transforms m/z data to neutral spectra both assumes and requires that multiple charge states *must* exist. Although this fundamental and universal observation has been described since the inception of electrospray, the reason for multiple protein m/z values is neither self-evident nor is it explained by present theories of electrospray ionisation. It does not happen during MALDI ionisation. Nor does it happen in solution, where chemistry assumes that all molecules of a protein under the same conditions possess the same charge (pI). Of the several mechanisms for electrospray ionisation proposed[8] [9], [10], [11] all address certain phenomena, but none provide a comprehensive explanation. It has been suggested that different mechanisms must exist to explain the differences observed between ionisation of small molecules and biopolymers, or those observed between native and denatured proteins [12]. Such explanations are intellectually unsatisfying, and a unified theory of electrospray is required.

## Results

We used four metrics to describe ESI spectra:

Maximum charge state (z_max_) was the highest observable charge state (or lowest m/z) at which ion intensity approached baseline.

Modal charge state (z_mode_) was the predominant species in a denaturing ion distribution.

Native charge state (z_native_) was the predominant species in a native ion distribution.

Full width half maximum of the native ion distribution (FWHM_native_) was measured in charge units.

### Charge and molecular flexibiliy

Electrospray charge states were determined for rigid self-assembled fibrils of diphenylalanine (FF) comprising of between 1 and 116 FF dipeptide units. (Fig1a). Each unit possessed a single m/z and the number of protons was directly proportional to fibril length. The discontinuous, stepped relationship occurs because of the unitary nature of charge. The number of protons acquired by unstructured oligomers of poly-L-lysine (Fig.1c) was also found to be proportional to polypeptide length, although in contrast to diphenyalanine, each possessed either one, two or three charge states, depending upon size. The larger oligomers carried not only higher charge, but also a greater number of m/z values. Next, proteins under native conditions were examined (Fig.1b). All of the 156 proteins displayed multiple m/z values. Small proteins had narrow charge distributions, becoming broader with increasing protein size. As with FF fibrils and polylysine, the absolute number of protons increased with size. Finally, denatured proteins and polypeptides were examined (Fig.1d). Again, all of the 212 proteins displayed multiple m/z values. While the smallest proteins had narrow charge distributions, these became proportionately broader with increasing linear sequence.

**Figure 1.**
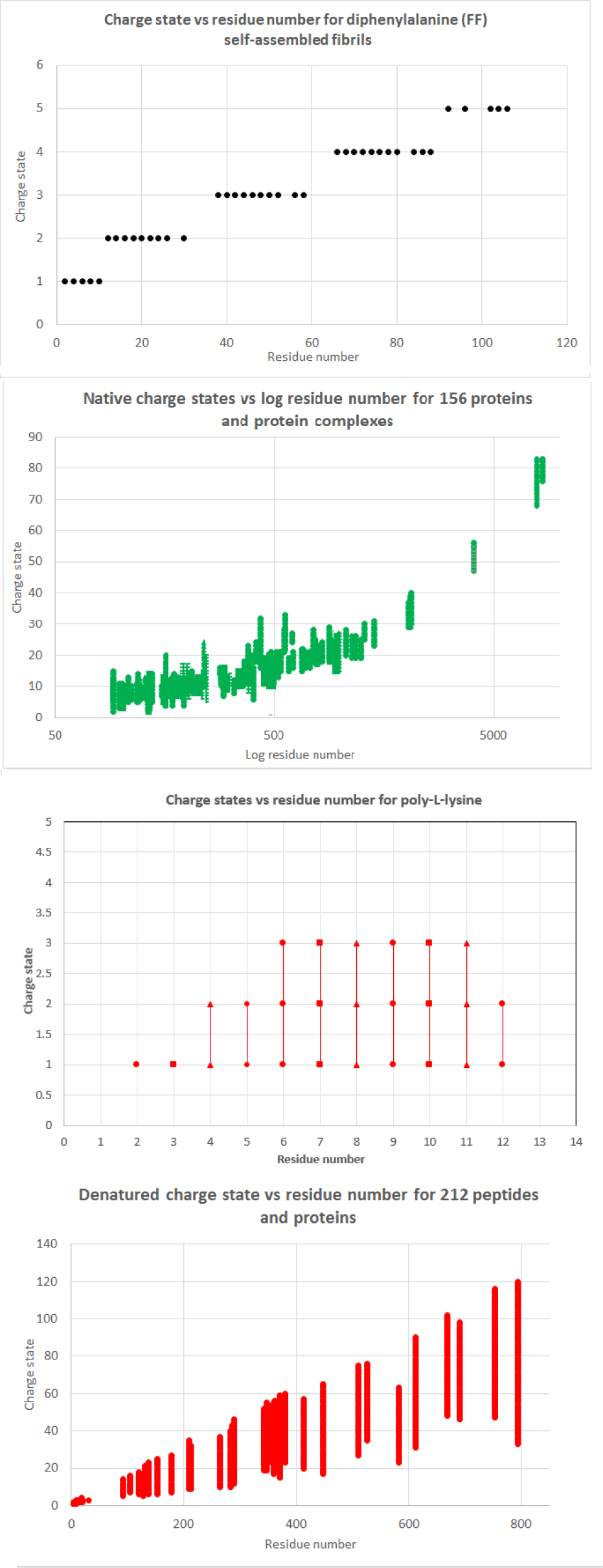
Relationship between charge and residue number for proteins and protein analogues with different degrees of flexibility: FF fibrils (a), native proteins (b), polylysine (c) and denatured proteins (d)

From figure 1d it may be seen that a linear relationship exists between minimum m/z (z_max_) and residue number or polypeptide chain length. z_max_ was plotted alone against residue number for 56 denatured proteins (fig.2) showing a highly significant correlation (R^2^ = 0.991). Eight proteins possessing intramolecular disulphide bonds were observed as outliers and were excluded from the initial regression calculation (Lysozyme, SLC1A4, β-lactoglobulin, PSF25, CD33, ELOVL7, KCNK10, BSA). Following reduction (and linearization) the z_max_ values of these proteins regressed to the trend line. No values appeared above the line.

**Figure 2.**
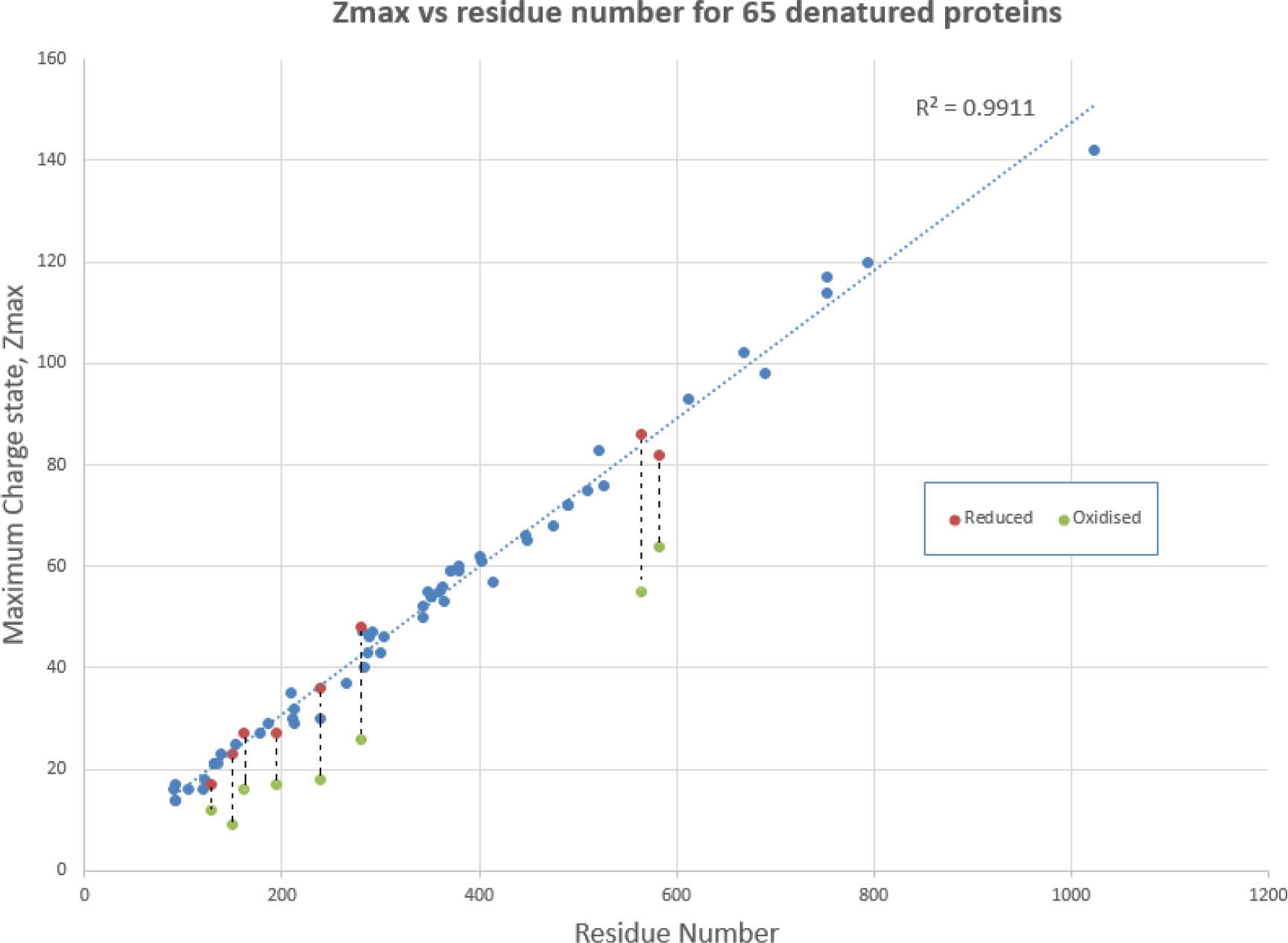
Relationship between maximum charge and residue number for denatured proteins. Proteins posessing disulphide bonds are shown in green. The same protein with reduced disulphide bonds is shown in red.

### Charge and solvent accessibility

Deuterium incorporation into ubiquitin was plotted against charge state under native conditions using published data from the original global exchange experiment of Katta *et al.* [13](Fig 3). Incorporation increased with increasing charge. An inflection point occurred between +8 and +9. For +7 and +8 deuterium incorporation was constant irrespective of charge, indicating that these m/z values possessed solvent accessibility to the same degree and the same conformation. This inflection point coincides with native/unfolded transition entropy barrier for ubiquitin.

**Figure 3.**
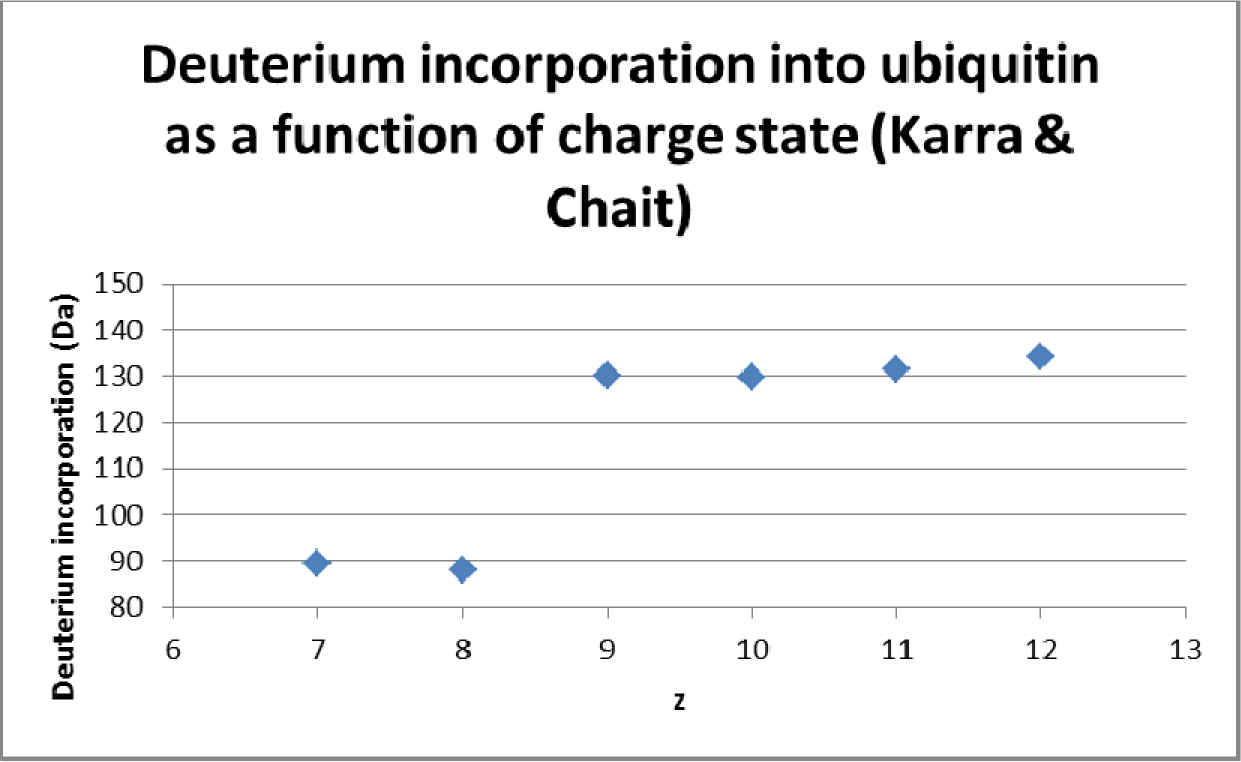
Relationship between deuterium incorporation and charge for native and denatured charge states of ubiquitin.

### Charge and disulphide bonds

Spectra were acquired for myoglobin and lysozyme under both denaturing (50 % methanol, 0.1 % formic acid, pH 2.0) and native conditions (50 mM ammonium acetate, pH 6.5). Myoglobin contains no disulphide bonds while lysozyme contains four. Native myoglobin displays a narrow distribution of four m/z values both with and without its bound haem cofactor (Fig 4a). Denatured myoglobin, displays a typical wide and skewed distribution of seventeen m/z values (Fig 4b). In contrast, spectra for lysozyme acquired under the same set of native and denaturing conditions shows narrow near symmetrical distributions of four or six m/z values which are almost identical and typical for native proteins (Figs 4c & 4d). Due to its four disulphide bonds, lysozyme is known to be stable under extreme conditions. Unlike myoglobin, both lysozyme spectra reflect native states and show that the m/z distribution here is dependent upon the degree of protein folding, rather than the electrospray solvent conditions.

**Figure 4.**
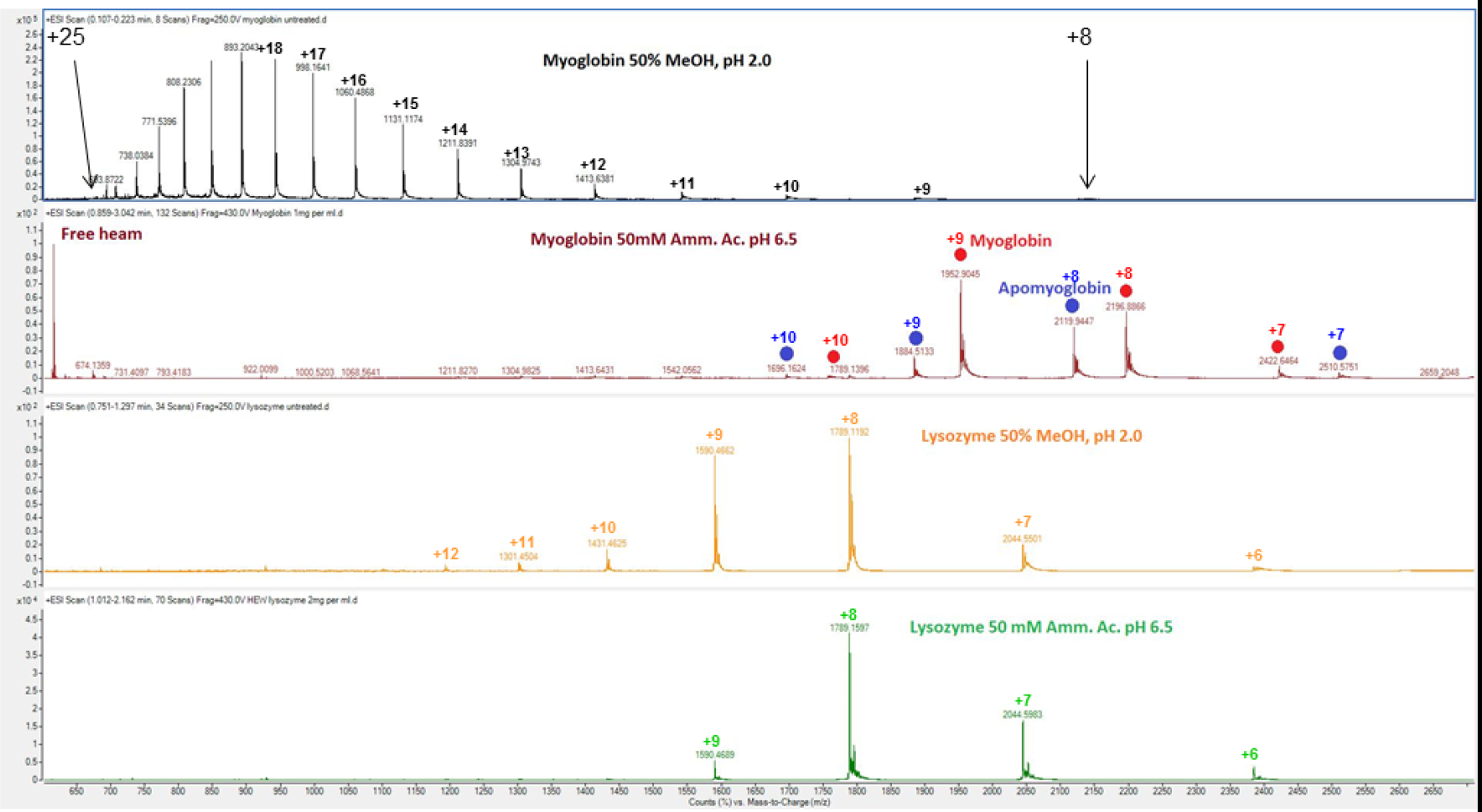
Effect of denaturing and native electrospray conditions upon the charge distributions of a typical protein, and a thermostable protein a) myoglobin denaturing b) myoglobin native c) lysozyme denatured d) lysozyme native.

Spectra were acquired for BSA under four conditions: native fresh preparation (Fig 5a), native aged preparation (Fig 5b) denatured with disulphides intact (Fig 5c) denatured with disulphides reduced (Fig 5d). An entropy barrier was observed at +20 marking the native/unfolded transition. The degree of protonation as well as the width of the m/z distribution was again dependent on the degree of unfolding, reaching maximum charge (z_max_) only when disulphides were reduced and the protein was able to achieve a near linear conformation.

**Figure 5.**
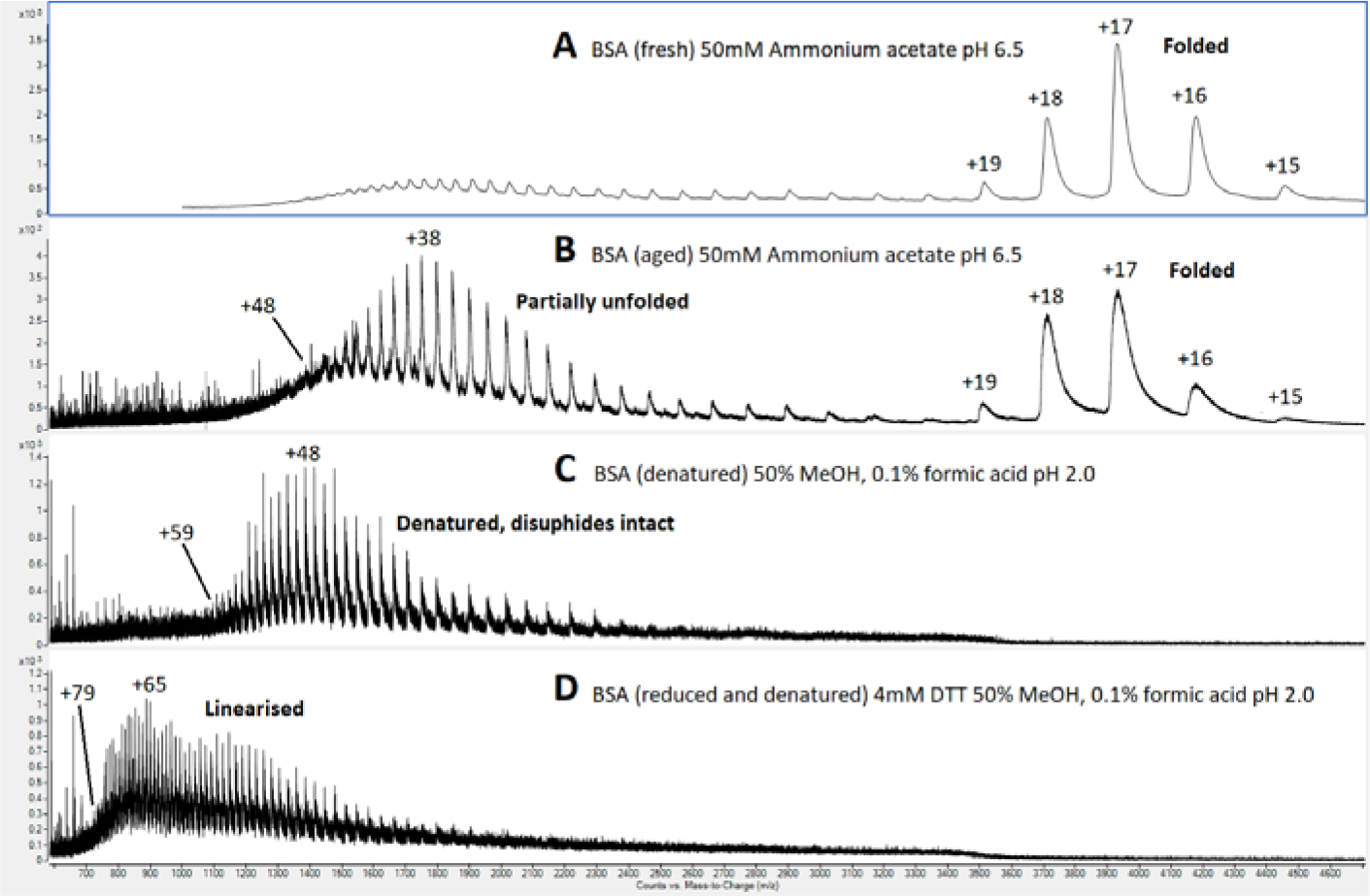
Effect of denaturation and disulphide bond reduction on the charge distribution of BSA a) native b) denatured, non-reducing c) denatured and reduced

### Charge and stability

Native spectra were acquired for trypsin under three conditions: i) native and enzymatically active, pH 8.0 ii) native, stable and enzymatically inactive, pH 1.0 iii) native, thermostabilized and active, pH 8.0. Using the calculation of Testa *et al.*[14] the radius for each m/z value was calculated and plotted against relative ion intensity. To allow comparison, peak height was normalised by division with the greatest value in each spectrum (Fig. 6). Narrow, symmetrical, native m/z distributions were observed in all cases. Significantly, the modal charge for thermostabilized or pH stabilized trypsin was one unit less, corresponding to a 1.5 Å reduction in diameter compared with active trypsin. In addition, both types of stablilized trypsin displayed narrower m/z distributions of 2.5 and 3.0 charge units full width at half maximum compared with 4.0 units for active trypsin at pH 8.0.

**Figure 6.**
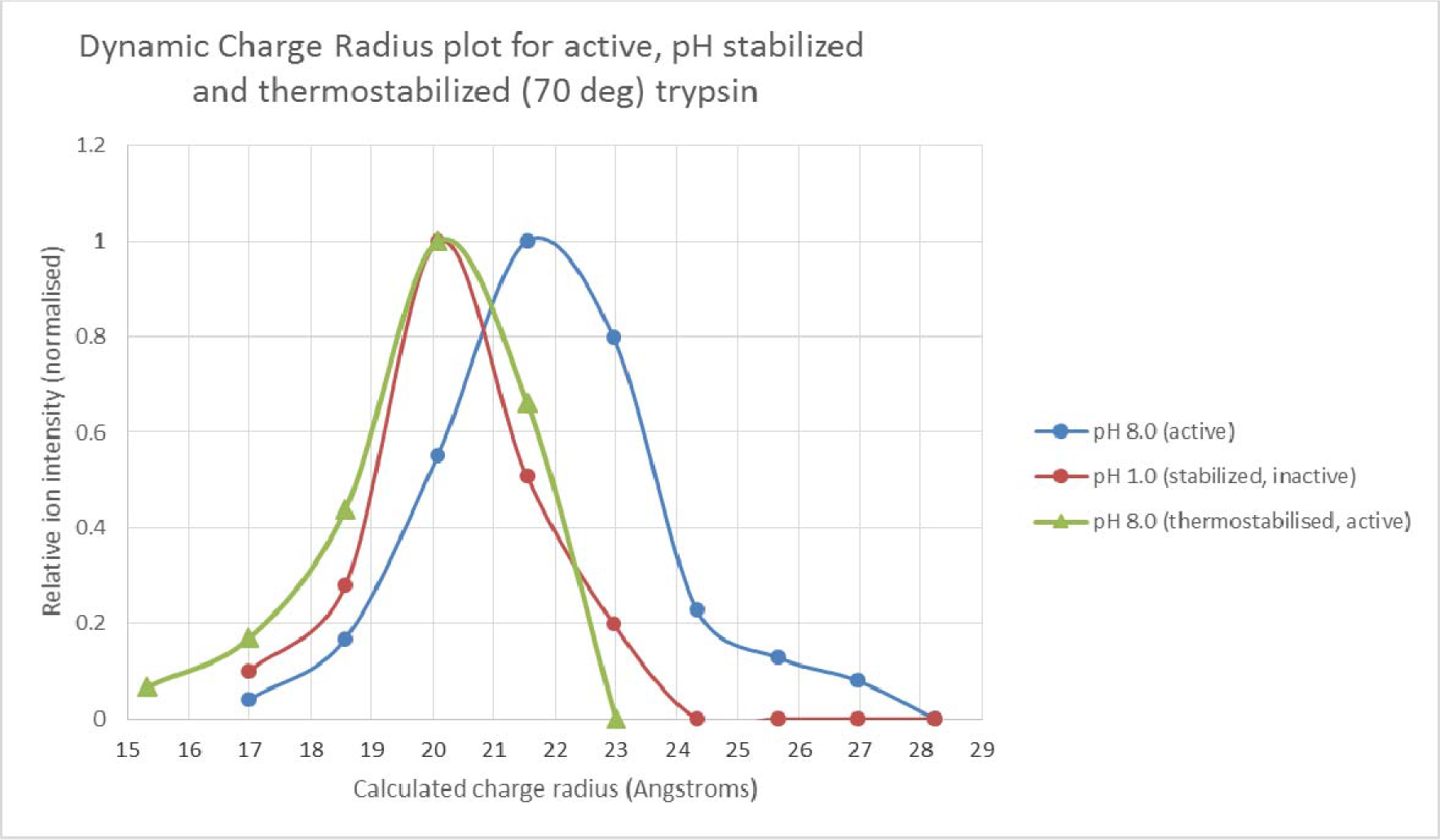
Effect of pH stabilization and thermo-stabilization upon calculated charge radius for an enzyme.

### Charge and phosphorylation

Spectra were acquired for the kinase Aurora B before and after autophosphorylation, generating phosphorylation states 0,1, 2, 3, 4 and 5. Using all m/z values for each phosphorylation state, the mean charge radius was calculated and plotted against phosphorylation state (Fig. 7). Radii for phosphorylation states 0,1 and 2 were calculated to be approximately 25.04 Å, while an inflection occurred at phosphorylation states 3, 4 and 5 corresponding to an increase of 0.12 Å. This is consistent with a charge induced conformational switch. Significantly (and counter-intuitively) the increase in negative charge in solution due to phosphorylation brings about an increase in *positive charge* following electrospray.

**Figure 7.**
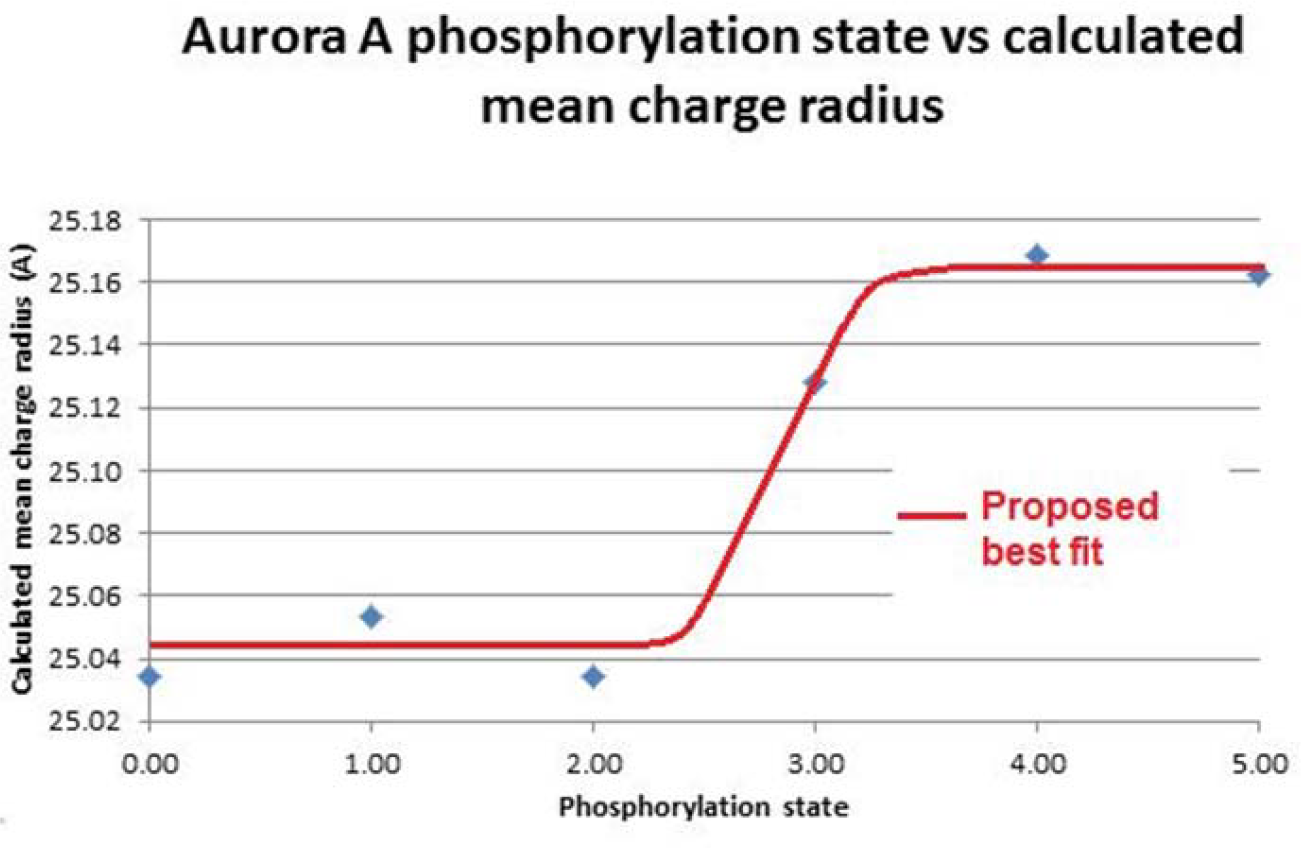
Effect of successive phosphorylation upon calculated charge radius for a kinase.

### Charge and ligand binding

Spectra were acquired for tetrameric enzyme HADH2 with and without bound cofactor NADH; and compared as described above for trypsin (Fig. 8). NADH binding results in a reduction in charge, equivalent to a 2.5 Å reduction in radius and consistent with adoption of a more compact conformation.

**Figure 8.**
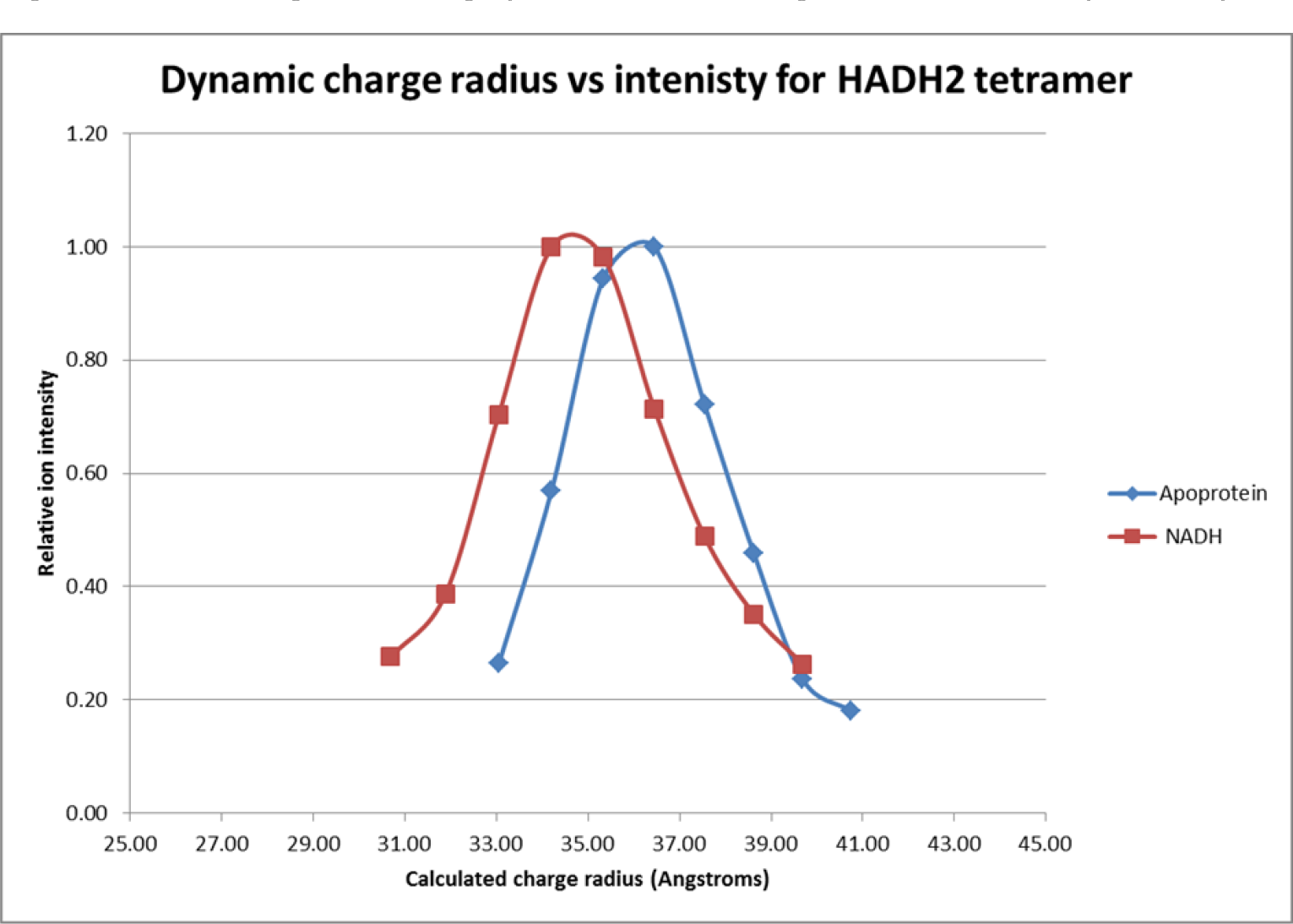
Effect of ligand binding upon calculated charge radius for an enzyme complex.

### Charge and pl

Spectra were acquired for pepsin and a 19-mer synthetic oligonucleotide, two highly electronegative species. Pepsin acquires 8-12 protons and 12-47 protons under native and denaturing conditions respectively (Figs. 9a & 9b), despite the fact that in solution it possesses only 5 basic functional groups. The oligonucleotide acquired 3 to 5 protons and possesses no basic functional groups (Fig.9c)

**Figure 9.**
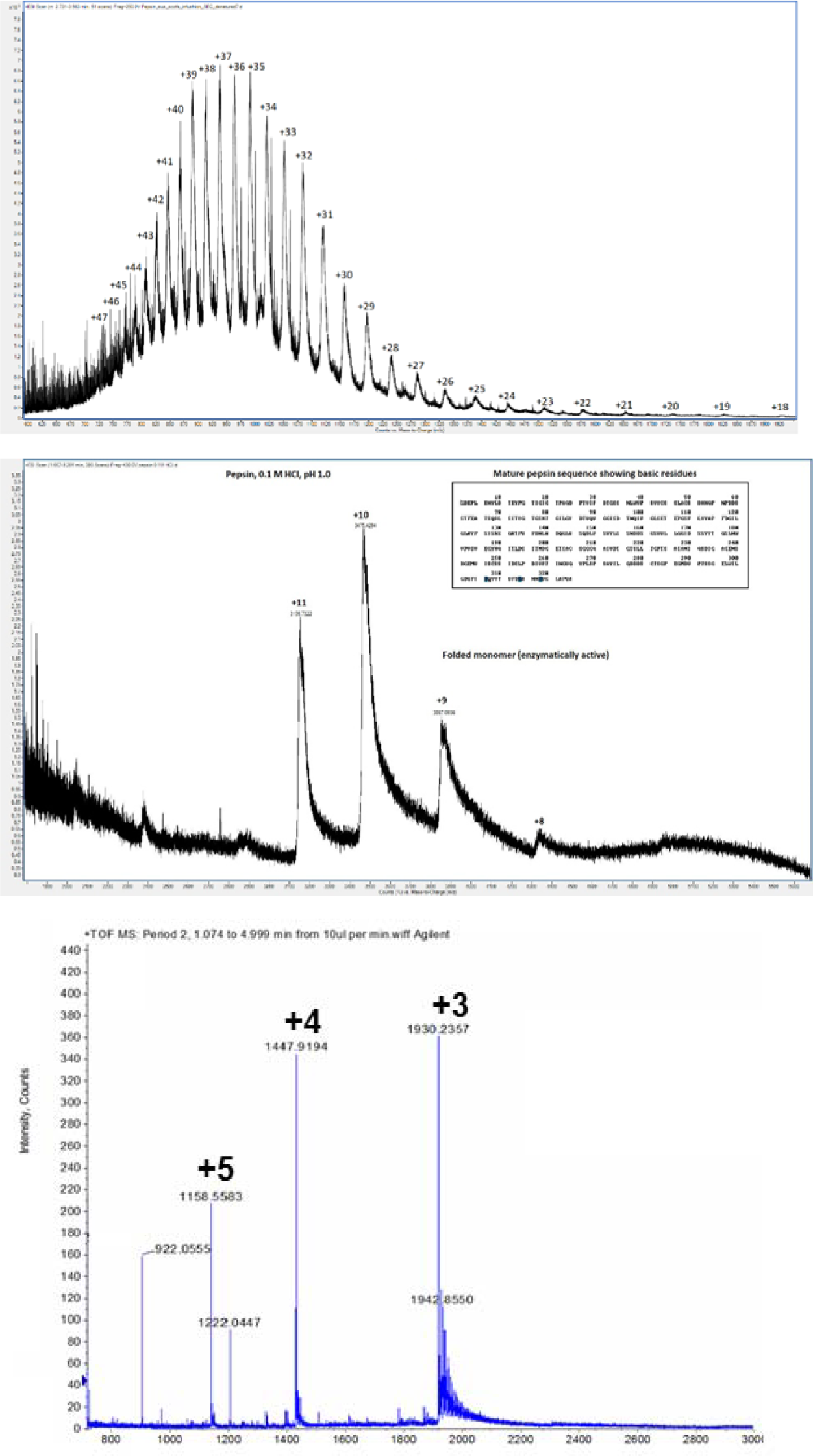
Electrospray protonation states for acidic biopolymers. a) denatured pepsin, pH 8.0 b) native pepsin, pH 1.0 c) 19-mer oligonucleotide, pH 6.5

Maximum ESI charge state (zmax) was determined for 25 representative proteins and this was subtracted from the total number of basic amino groups available for chemical ionisation in solution. The resulting plot shows that nearly all proteins are protonated in electrospray to an extent far beyond that which is possible in solution. Acidic proteins such as pepsin display the largest difference, whereas those proteins such as BSA which have a lower ESI protonation state display multiple disulphide bonds.

### Charge and size

The protein KCTD17 is known from both crystallography and SAXS [15] to form a 924,253 Da perfect hollow sphere composed of 60 identical subunits and 84 Å in diameter. Two spectra for the complex were acquired at different protein concentrations yielding modal charge states of +78 and +79 corresponding to radii of 85.5 Å and 83.5 Å in agreement with the crystallography measurements (Fig 10). Furthermore, these measurements appear to show that charge accumulates only on the *outer* surface of the KCTD17 sphere.

**Figure 10.**
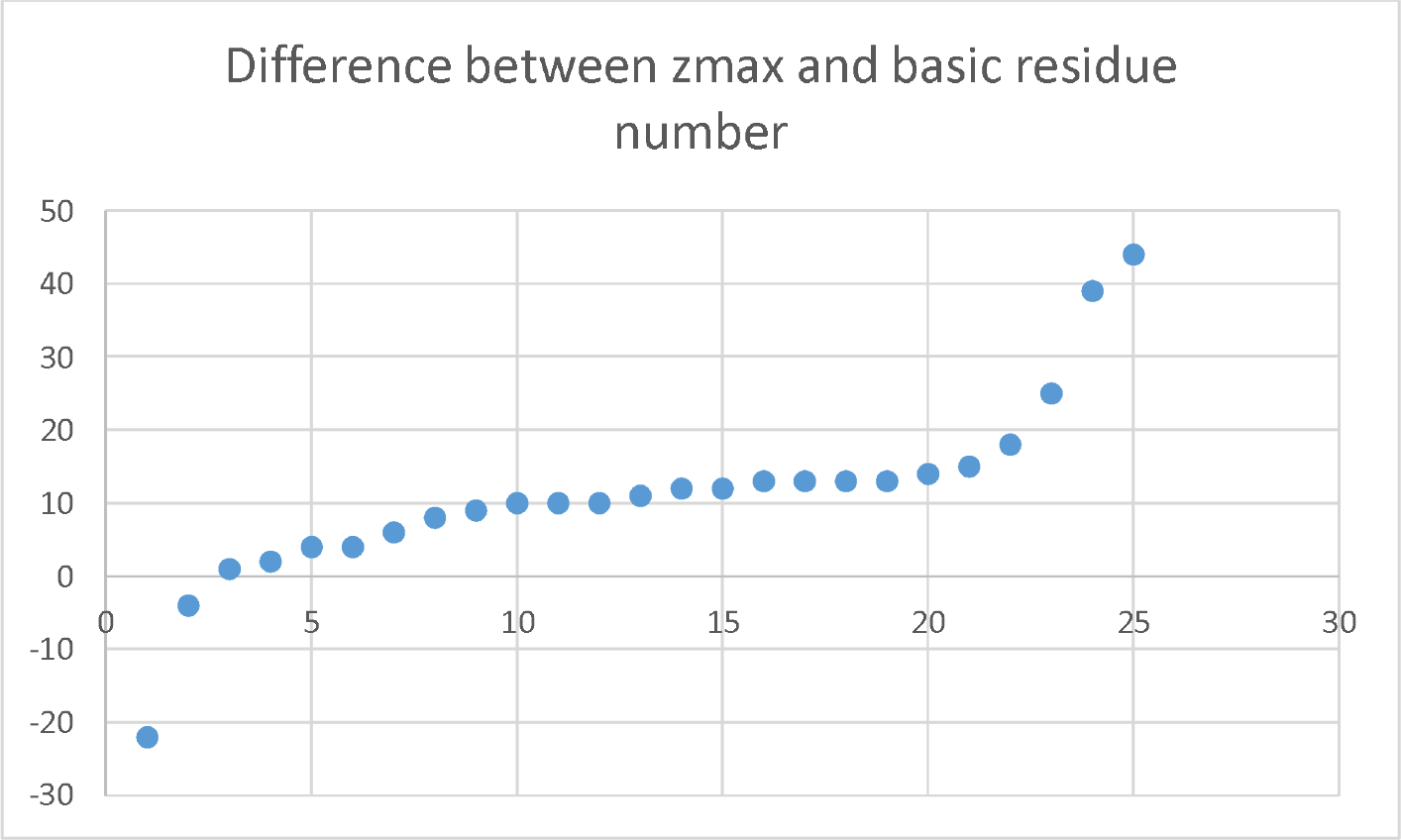
Difference between maximum charge state and basic residue number for 25 representative proteins

### Charge and shape

Spectra were acquired for the developmental regulator protein Sonic Hedgehog which is known to form a homodimeric complex (Fig 11). Unusually, the m/z values for the monomer (m1 z8, m1 z9 & m1 z10) were found to overlap completely with those of the dimer (m2 z17, m2 z18 & m2 z 19). This overlap is possible only if the dimer carries exactly double the charge of the monomer and possesses twice its surface area. This, in turn, is possible only if the dimer subunits share a single point of contact. Crystallographic studies of the Shh dimer[16] show that this is indeed the case.

**Figure 11.**
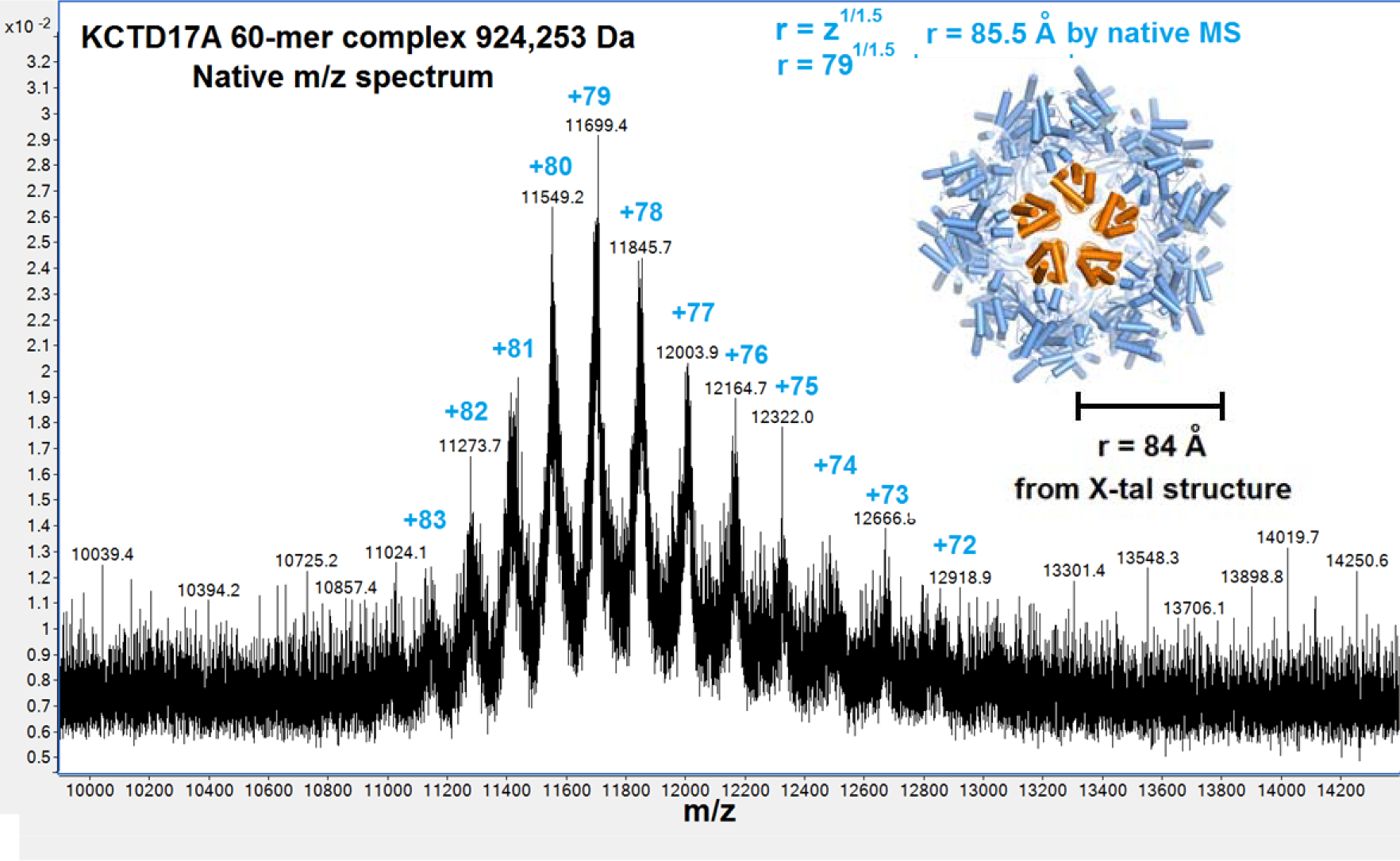
Charge radius measurement for a spherical protein complex

### Charge and fragmentation

Chymotryptic digests of BSA were analysed by LC-MSMS with selection of +1, +2 and +3 parent ions. Singly charged ions AVSVLL+ and FYAPELL+ were found to generate both b and y fragment ions indicating that the single proton is able to generate charged fragments from both C and N termini (Fig 12). This strongly suggests that under these conditions electrospray protons are able to exist in more than one location.

**Figure 12.**
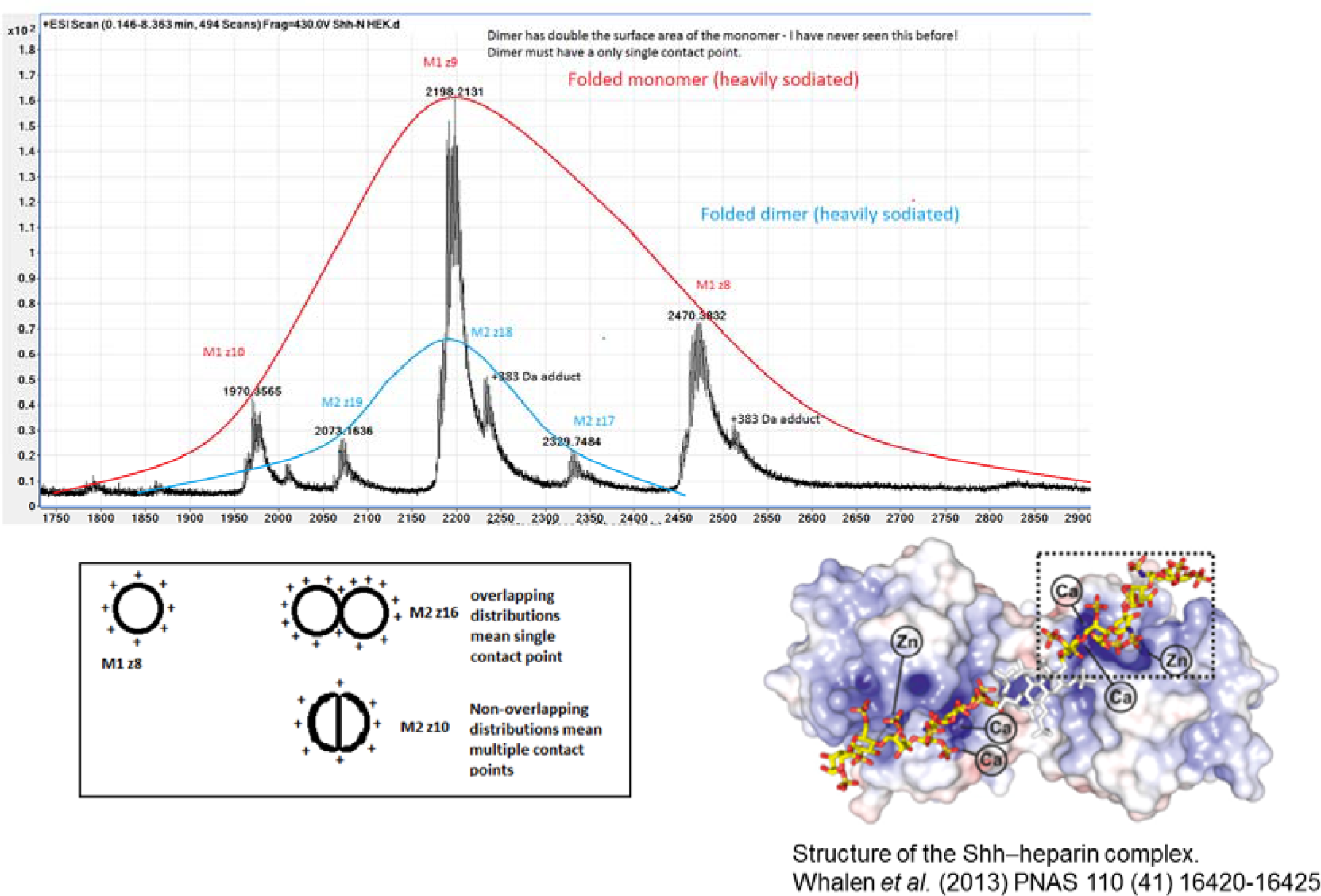
Inference of Sonic hedgehog dimer shape from charge/surface

### Charge and free energy

The Thomson Problem describes the minimum free energy of charged particles obeying Coulomb’s law on the surface of a sphere [17]. We plotted the minimum free energy for n charged particles against the charge radius for the same number of charges using the equation z=r^1.5^ [14] and observed a perfect correlation (Fig. 13)

**Figure 13.**
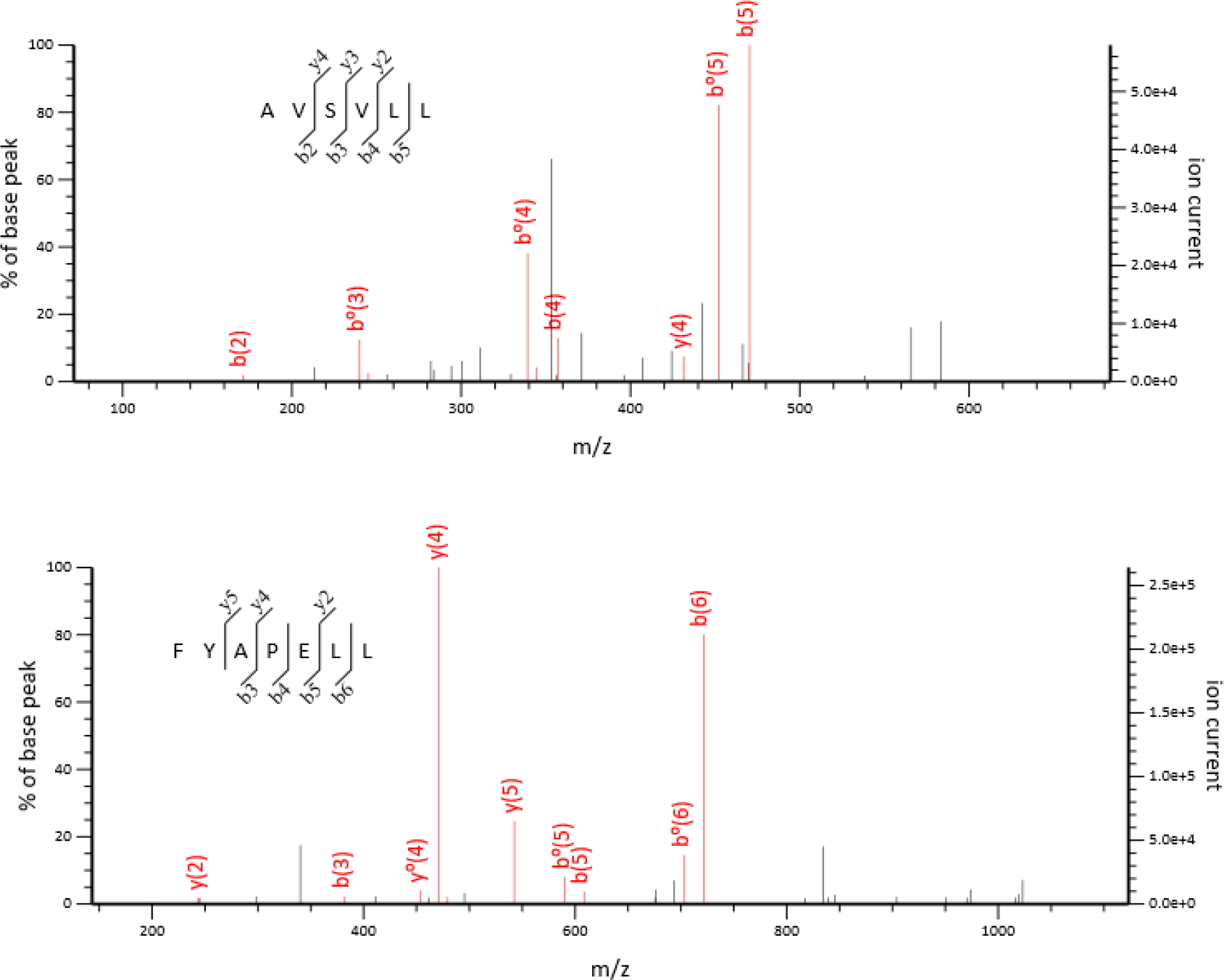
Mascot fragmentation data showing both N and C terminal protonated ions derived from the same singly charged peptides

**Figure 14.**
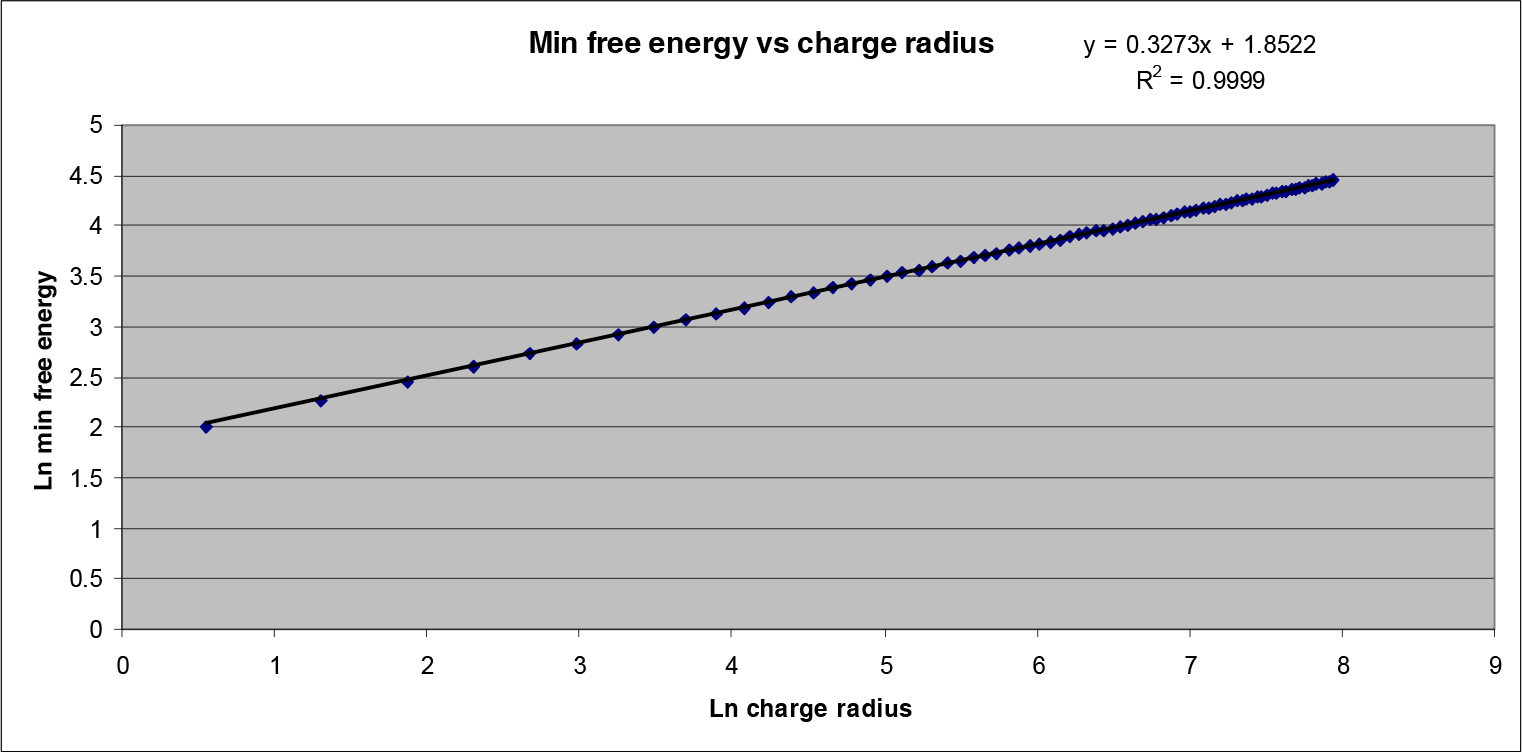
Plot of Thomson free energy against charge radius for a spherical protein

## Discussion

Native spectra can only be interpreted correctly if the mechanism determining charge and intensity for each ion is understood. The universal observation of multiple protein m/z values requires an explanation, which current models for electrospray ionisation do not provide [8, 9,12]. Figures 5, 7 and 9 lead us to conclude that m/z cannot be dependent upon charge in solution. The higher charge states for myoglobin and BSA do not arise directly from low pH, but indirectly as a result of unfolding. Similarly, so-called supercharging reagents play no direct role in the ionisation process, but are mild denaturants creating structures with larger surface area[18]. Kaltashov and Mohimen [19] established a definitive linear relationship between native charge state and surface area. This should underpin our understanding of the electrospray process, and other observations need to be consistent with it. Towards this goal, Testa *et al.* demonstrated an approximate relationship between modal charge and surface area for denatured proteins[14]. In this paper we chose a different metric, maximum charge, and demonstrate a definitive linear relationship with surface area for denatured proteins. Furthermore, we show that if the linear maximum surface area conformation is prevented by intramolecular disulphide bonds, maximum charge state is not reached. Orthogonal observations by Rinke *et al*.[20] using ion beam deposition mass spectrometry coupled to superconducting tunnelling microscopy showed that charge state for denatured chytochrome C is related to the degree of unfolding, and that linearised protein carries the maximum charge. Although the argument that unfolded proteins expose a greater number of charge carrying amino acid side chains is plausible, electrospray ions of pepsin, for example, bearing 34 protons yet only 5 protonated side chains demand an alternative explanation, as pointed out previously[21]. Pepsin is highly acidic and atypical, but most proteins carry a greater positive charge during electrospray than is possible in solution as illustrated in figure 10. It is incumbent upon any model to explain not just the presence but also the relative abundance of all m/z species. Our data implies that under all conditions charge is dependent upon surface area: surface area is determined by protein folding. Maximum charge density for proteins is never exceeded and corresponds to the extended linear conformation.

We used charge to measure the diameter of a perfectly spherical protein complex of KCTD17, and determined by crystallography that this measurement is accurate to within 2%. Crystallography and SAXS data both indicate that the complex is a hollow sphere with two solvent accessible surfaces [15]. From agreement with the calculation of Testa *et al*.[14], we conclude that charge is present on the outer surface only. We have used charge to measure conformational changes in surface area resulting from thermostabilization, pH stabilization, phosphorylation and ligand binding. The resulting changes in surface area are as expected and the same degree of accuracy is assumed.

From figures 1 and 5 we conclude that the number of m/z values is for a species is related to its flexibility. Single m/z values for FF fibrils are highly significant in the light of our other observations. FF spontaneously forms self-assembled fibrils[22]. These were of interest because chemically, they are analogous to polypeptide, yet unlike polypeptides they form a rigid structure. Each fibril possesses a single m/z in contrast with polylysine, a flexible homopolypeptide of varying length possessing one, two or three m/z values. Also in contrast to FF, proteins possess multiple m/z values under all conditions. Larger proteins have wider ion distributions, denatured proteins always having substantially greater distributions than their native counterpart. If multiple m/z values are a measure of flexibility, they should not be observed in rigid structures. Conversely, flexible structures should not have a single m/z. Our data shows that this is the case. Moreover, the degree of flexibility observed for native proteins is in the region of 10 Å and consistent with collective molecular motion.

It follows from these observations that m/z species in a native ion distribution do not arise from different molecular structures. Rather they arise from a single, stable native conformation undergoing molecular motion. Hydrogen-deuterium exchange experiments indicate that incorporation is constant for these distributions. Similarly, the linear relationship between ion mobility drift time and charge for native m/z distributions indicates that in the gas phase these ions have the same collisional cross sectional area.

We explain these observations in terms of electrostatic charging, and propose the following model of positive electrospray ionisation for species above the Rayleigh limit[23]:

1. An excess of protons is always available.
2. Protons are mobile over the molecular surface and obey Coulomb’s law to achieve minimum free energy.

From this simple model, it follows that the maximum charge state corresponds to the largest possible surface area, which for proteins is a linearized polypeptide chain; and that the minimum charge state corresponds to the smallest possible surface area, which for proteins, is usually the native conformation. For a native globular protein the radius may be approximated by z 1/1.5 [14].

The narrow Gaussian distribution of m/z peaks always observed for native proteins is the result of dynamic changes in surface area resulting from restricted, defined molecular motion as the protein is ionised and enters the gas phase. Denatured proteins undergo random molecular motion and adopt any number of unfolded states. These too form a Gaussian distribution of m/z peaks which is skewed towards a linear conformation of maximum charge density. The abundance of any m/z is proportional to the number of random conformers sharing the same surface area. Since linearization and maximum charge is an energetically unfavourable minimum entropy state, this m/z will be low in abundance or not present at all. In contrast, the most abundant m/z represents the surface area possessed by the greatest number of random conformers and is thus the maximum entropy state for a denatured protein.

Although mobile protons would be expected to obey Coulomb’s law, and this model is in agreement with experimental observations, evidence for mobile protons is needed. We observed in figure 13 that fragment ions can be generated from both ends of a singly charged peptide, meaning electrospray protons are not fixed in one location. The mobile proton theory is not new[24],[25]. Transitory excitation of protons becoming mobile during CID is used to explain peptide fragmentation. We propose that protons are in fact mobile throughout the gas phase ion stage. There are few NMR studies of proteins in the gas phase but, Oomens [26] reported the presence of a 1483 cm^-1^ band from cytochrome c. This was absent from the liquid phase and appeared to be charge state dependent. The same charge state-dependent phenomenon has been observed for ß-lactoglobulin[27]. We propose that 1483 cm^-1^ corresponds to the mobile proton shell for these proteins. In the liquid phase, protons must reside at basic functional groups. However, commonplace electrostatic charging induced by friction between two insulators has no requirement for this. Charges are trapped on the surface, and in an analogous fashion, protons are trapped on the surface of an ion by the surrounding vacuum.

In addition to experimental evidence, a theoretical basis for our model is also required. The behaviour of multiple charges constrained to the surface of a sphere was first considered by JJ Thomson in his early model of atomic theory[17]. While the model has been superseded, the Thomson Problem was identified as one of eighteen unsolved problems in mathematics [28]. It defines the spatial arrangement for a given number of charges necessary to achieve the lowest possible free energy while obeying Coulomb’s law (which states that the force between two equal charges is inversely proportional to the square of the distance between them [29]). Our theoretical calculation of charge radius on a sphere demonstrates perfect correlation with Thompson free energy. As the father of mass spectrometry, it is highly appropriate that JJ Thomson’s work remains relevant.

Native mass spectrometry is only of value if m/z measurements truly reflect conditions in the liquid phase. Ion mobility describes the behaviour of ions exclusively in the gas phase. This may or may not reflect prior conditions in solution. Use of the term “near native” [30] is an indication of this uncertainty, but defies scientific definition. Proteins in the native state must retain all characteristics which define that state: activity, stability, conformation, flexibility and molecular interaction. An entropy barrier must be overcome to reach a non-native partially or fully unfolded state, hence proteins are either native or they are not. The distinction is obvious from the biphasic ion distributions of m/z spectra such as that for BSA in figure 5b, and no alternative explanation for this has ever been proposed. While measurement of m/z occurs in the gas phase, *assignment of charge*, namely ionisation itself, cannot. Transfer of charge from the ESI droplet surface to the analyte within it must have already occurred when solvent evaporation is complete. Thus ionisation occurs at or prior to the solution phase/gas phase transition point. Once in the gas phase, conformation may alter, but the mass to charge ratio remains fixed and thus reflects the condition of the protein in solution immediately prior to ionisation. Being a gas phase only phenomenon, ion mobility is a distraction from understanding conformation in solution. If m/z distributions display *all* the characteristics expected of native proteins in solution, including molecular motion as we have shown, then logically they must represent that state.

We have demonstrated that the relationship between ESI charge and protein surface area is universal, and that multiple charge states are dependent upon the degree of molecular motion. The electrostatic model proposed is consistent with our observations and is predicted by established principles of physics. Independent evidence from hydrogen-deuterium exchange, ion beam deposition, peptide fragmentation and infrared NMR supports this model. Surface area and critical alterations of it resulting from conformational change can be measured with greater precision than other biophysical methods, and since charge state is an intrinsic ion property, calibration is not required. Charge state distribution analysis is thus a simple and powerful tool to explore protein structure in solution.

## Materials and Methods

Diphenylalanine fibrils were prepared as follows. FF (Sigma) was dissolved in water at a concentration of 2 mg/ml by heating to 65 °C and subsequently allowed to cool to ambient temperature. Polylysine (Sigma) was prepared by dilution in water at 1 mg/ml. Analysis was by direct infusion at 6 μL/min. Charge state for each species was determined by observed separation between ^12^C and ^13^C isotopes. Myoglobin, lysozyme, BSA, beta-galactosidase, pepsin and proteomic grade trypsin were obtained commercially (Sigma). “Smart Digest” thermostabilised trypsin was a gift from the Thermo Corporation. Sonic Hedgehog was expressed and purified as described elsewhere (Born). KCTD17, Aurora kinase, HADH2 and the remaining proteins used in this study were cloned, expressed and purified in-house following protein production protocols deposited with the relevant PDB structure or freely available in accordance with the SGC open-access research policy (http://www.thesgc.org/scientists/proteins). Disulphide bond reduction was achieved by addition of DTT to 1 mM and incubation for 20 minutes at room temperature. Aurora kinase phosphorylation was achieved by incubation of protein at 1 mg/ml with 1mM ATP and 2 mM MgCl_2_ overnight at 4 °C.

Proteins for native mass spectrometry were buffer exchanged into 50mM ammonium acetate by 3 rounds of gel filtration using BioGel P6 (Biorad) spin columns according to the manufacturer’s instructions. All intact mass spectra were acquired using an Agilent 6530 QTOF operating in positive ion 2 GHz mode up to m/z 3,500 or 1 GHz mode up to m/z 20,000. Samples were introduced via an Agilent 1290 HPLC under denaturing conditions or via a syringe pump under native conditions. Denaturing mass spectra and native spectra were acquired as described previously [7] [31]. LC-MSMS spectra were acquired on a Bruker HCT ion trap coupled to a Thermo U3000 nano HPLC using previously described protocols [7],

